# TNMplot.com: a web tool for the comparison of gene expression in normal, tumor and metastatic tissues

**DOI:** 10.1101/2020.11.10.376228

**Authors:** Áron Bartha, Balázs Győrffy

**Author notes:** **Correspondence:** Balázs Győrffy MD PhD DSc, Department of Bioinformatics, Semmelweis University, Tűzoltó u. 7-9, 1094, Budapest, Hungary.

## Abstract

Genes showing higher expression in either tumor or metastatic tissues can help in better understanding tumor formation, and can serve as biomarkers of progression or as therapy targets with minimal off-target effects. Our goal was to establish an integrated database using available transcriptome-level datasets and to create a web-platform enabling mining of this database by comparing normal, tumor and metastatic data across all genes in real time.

We utilized data generated by either gene arrays or RNA-seq. Gene array data were manually selected from NCBI-GEO. RNA sequencing data was downloaded from the TCGA, TARGET, and GTEx repositories. TCGA and TARGET contain predominantly tumor and metastatic samples from adult and pediatric patients, while GTEx samples are from healthy tissues. Statistical significance was computed using Mann-Whitney or Kruskall-Wallis tests.

The entire database contains 56,938 samples including 33,520 samples from 3,180 gene chip-based studies (453 metastatic, 29,376 tumorous and 3,691 normal samples), 11,010 samples from TCGA (394 metastatic, 9,886 tumorous and 730 normal), 1,193 samples from TARGET (1 metastatic, 1,180 tumor, 12 normal) and 11,215 normal samples from GTEx. The most consistently up-regulated genes across multiple tumor types were TOP2A (mean FC=7.8), SPP1 (FC=7.0) and CENPA (FC=6.03) and the most consistently down-regulated gene was ADH1B (mean FC=0.15). Validation of differential expression using equally sized training and test sets confirmed reliability of the database in breast, colon, and lung cancer (p<0.0001). The online analysis platform enables unrestricted mining of the database and is accessible at www.tnmplot.com.

## INTRODUCTION

Cancer emerges as normal cells mutate first to pre-cancerous, then to malignant cells because of genetic or epigenetic lesions. Such lesions originate mostly in external mutagenic factors but hereditary mutations also influence the evolution. These genetic lesions lead to gene expression changes in the tumor cells which gear up the cancerous phenotype [1].

While most genes exhibit comparable expression profiles between cancerous and normal tissues, those differentially expressed can serve as either targets of treatment or molecular biomarkers of cancer progression. Targeting a gene with higher expression of a certain gene product can deliver astonishing clinical benefit as was demonstrated over two decades ago by the selective inhibition of overexpressed tyrosin kinases [2].

Gene expression changes in cancer cells are related to a limited set of special characteristics often termed as cancer hallmarks [3]. These paramount differences between malignant and normal tissues include among others resistance to cell death and activating invasion and metastasis. Various experimental methods capable of inspecting these hallmark genes have been reviewed previously [4]. Currently, the most widespread and robust techniques to determine transcriptome-level gene expression include RNA-sequencing and microarray platforms, while selected genes can be measured by RT-qPCR or NanoString technologies [5].

Both RNA-seq and microarray techniques produce a vast amount of clinically relevant data and large repositories hosting thousands of samples are now available. The National Cancer Institute’s Genomic Data Commons (GDC) platform provides whole exome sequencing data and transcriptome level gene expression datasets such as The Cancer Genome Atlas (TCGA)[6] and the Therapeutically Applicable Research to Generate Effective Treatments (TARGET)[7]. The Genotype-Tissue Expression (GTEx) repository makes available RNA sequencing, exome sequencing and whole genomic data for the same patient [8]. Nevertheless, the largest open resource is the Gene Expression Omnibus of National Center for Biotechnology Information (NCBI-GEO), which provides microarray, next-generation sequencing and additional high-throughput genomics data for hundreds of thousands of samples [9]. A common feature of these repositories is the provision of raw data in addition to processed and aggregated results.

At the same time, digesting such large sample cohorts requires complex bioinformatical analytical tools and it can be also time-consuming. Mining these databases could be speeded up by an openly available, validated and easily accessible online tool which enables the comparison of expression profiles between normal and cancer related data. Our first aim was to establish an integrated database of a significant number of normal and tumor samples with transcriptome-level gene expression data. We sought to establish a database which includes both adult and pediatric cases and both RNA-seq and gene array datasets. Our second goal was to validate the reliability of the database by employing a training-test approach to identify genes showing differential expression in selected tumor types. Finally, we designed an online analysis portal which can enable the comparison of gene expression changes across all genes and multiple platforms by mining the entire integrated database.

## MATERIALS AND METHODS

### Database setup – gene arrays

We searched the NCBI Gene Expression Omnibus (https://www.ncbi.nlm.nih.gov/geo/) repository for datasets containing “cancer” samples. Only datasets utilizing the Affymetrix HGU133, HGU133A_2 and HGU133A platforms were considered because these platforms use identical sequences for the detection of the same gene. In total 3,180 GEO series met these criteria, and each of these has been manually examined. We executed a filtering to exclude datasets containing either cell line studies, pooled samples, or xenograft experiments. Samples taken after neoadjuvant therapy were also excluded. In addition, samples with incomplete description, unavailable raw data and repeatedly published samples with distinct identifiers have been removed. For this, the expression of the first 20 genes were compared, and samples with identical values were identified. In each case, the first published version was retained in the dataset. Following this manual selection, the remaining samples were normalized using the MAS5 algorithm by employing the Affy Bioconductor library [10]. Finally, a second scaling normalization was made to set the mean expression on each array to 1000.

### Database setup – RNAseq

RNAseq data for a total of 11,688 samples were downloaded from the Genotype-Tissue Expression (GTEx) portal (version no. 7), from which two non-primary cohorts have been removed. Cell line studies available in GTEx were omitted. Read counts were normalized by the DESeq2 algorithm [11], followed by a second scaling normalization. Using the GDC database’s (https://portal.gdc.cancer.gov/) TCGA and TARGET projects, 11,010 and 1,197 files were downloaded, respectively. We only included primary tumors, adjacent normal, and metastatic tissues. Thus, non-primary tissue samples have been excluded. HTSeq – Counts files were normalized by DeSeq2 and a second scaling normalization was also executed for both cohorts.

### Gene annotation

In order to select the optimal probe set for each gene, we used the JetSet [12] correction and annotation package which delivered 12,210 unique genes in the gene-array datasets. Appropriate genes in the RNA-seq cohorts were selected and annotated by the biomaRt [13] and AnnotationDbi [14] R packages. The number of unique genes remaining after gene selection in the GTEx, TARGET, and TCGA databases was 21,479. After harmonization the GTEx and GDC data were combined into a single set. For the support of future data analysis, we constructed a master gene annotation data table with all the previous gene names and available synonyms for each included gene (**Supplemental Table 1**).

### Statistical analysis

Data processing and analysis features of the TNM-plotter pipeline were developed in R version 3.6.1. Comparison of the normal and the tumorous samples was performed by Mann-Whitney U test, matched tissues with adjacent samples were compared using the Wilcoxon test. Normal, tumorous and metastatic tissue gene comparison can be analyzed using Kruskal-Wallis test. Statistical significance cutoff was set at p<0.01.

### Shiny user interface

Graphical visualization including box plots, bar charts, and violin plots produced by the TNM-plotter algorithm were developed using the ggplot2 R package[15]. The web application and the user interface was developed by employing Shiny R packages, with the utilization of the ShinyThemes (http://rstudio.github.io/shinythemes/) and the ShinyCssLoaders (https://github.com/daattali/shinycssloaders) R packages [16].

### Validation of differential expression

In order to validate the effectiveness of the proposed approach and to confirm the reliability of the integrated database, we conducted a validation using randomly selected training and test sets across breast, lung and colon tissue dataset in both RNA-Seq and gene array platforms. In this validation process we compared the expression profiles of normal and tumor samples using the Mann-Whitney U test for 12,210 genes in the GEO and for 21,479 genes in the GDC datasets. Following calculation of the p values for each gene, a Chi-squared test was performed to compare selection overlap between the training set and the test sets. Volcano plots comparing −log_10_ p values and Log2 fold changes were generated to visualize differential expression.

### Cancer biomarker genes

To pinpoint genes showing the highest differential expression between normal and tumor samples across multiple tumor types we utilized the analysis pipeline and the database of the top ten cancer types with the highest mortality. Tumor types were selected using the 2019 mortality data from the United States [17]. We compared gene expression values between normal and tumor samples for all available genes in all platforms in each selected tumor type using the Mann-Whitney U test. Then, to combat multiple hypothesis testing we calculated the False Discovery Rate using the Benjamini-Hochberg method. Subsequently, the remaining significant genes were ranked by using the median fold change (FC) in all tissues. In other words, the significant genes were ranked based on their gene expression differences across all investigated tumor types. Finally, we selected genes with the highest FC values in both RNA-seq and gene array datasets.

## RESULTS

### Integrated database

The entire database holds altogether 56,938 samples including both RNA-seq and gene array samples.

These include after pre-processing 33,520 unique gene array samples from 38 tissue types, including 3,691 normal, 29,376 tumorous and 453 metastatic samples. For each of these samples, the mRNA expression of 12,210 genes is available.

Included RNA-seq data comprise three different platforms. After curation, normalization steps and data processing we collected data of 11,010 samples including 730 normal, 9,886 cancerous and 394 metastatic specimens from adult cancer patients. We also added 1,193 pediatric related data from GDC consisting of 12 normal, 1,180 cancerous, and 1 metastatic samples. In order to increase the number of normal samples we included further 11,215 RNA-Seq GTEx data form non-cancerous persons. Steps of data curation and processing are summarized in **Table 1**.

**Table 1.**
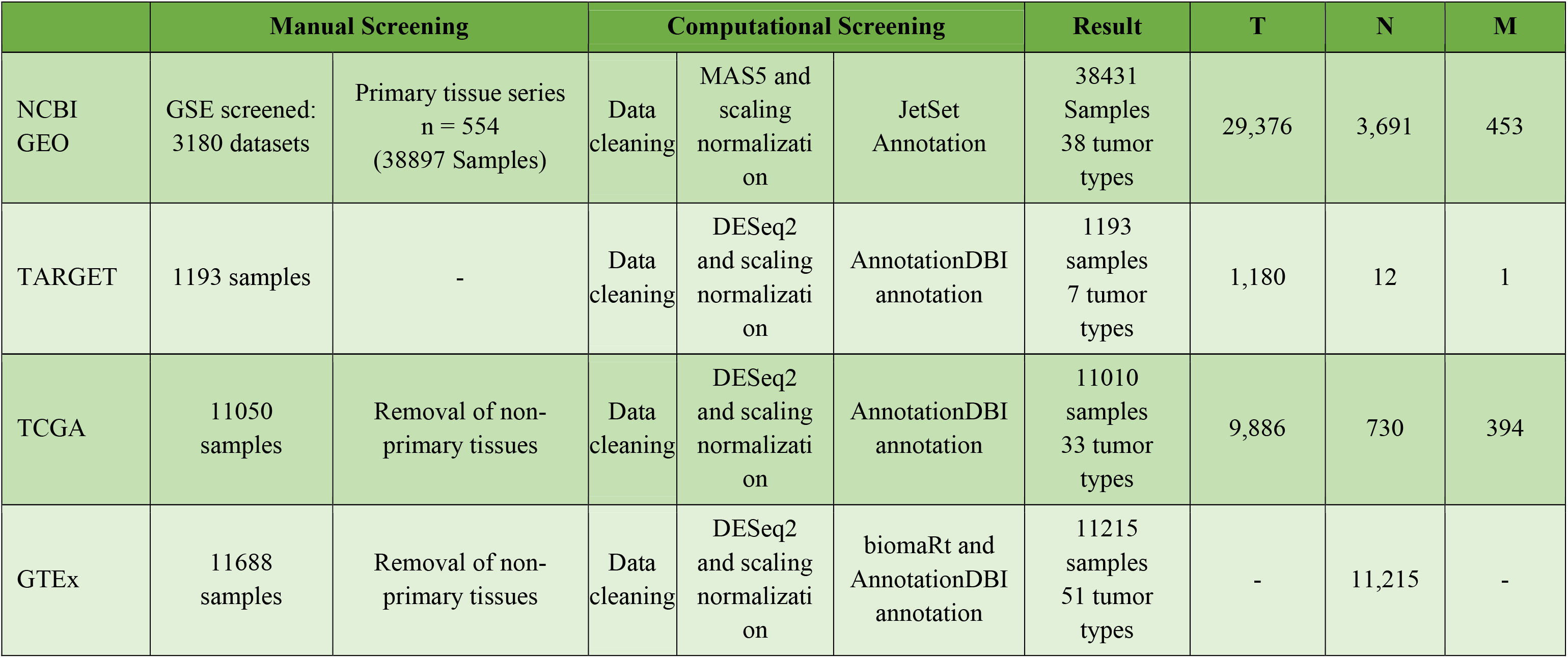
Summary of datasets and data processing.

### TNMplot.com analysis platform

We established a web application to enable real-time comparison of gene expression changes between tumor, normal and metastatic tissues amongst different types of platforms across all genes. The registration-free analysis portal can be accessed at www.tnmplot.com and has three separate analysis options. The pan-cancer analysis tool compares normal and tumorous samples across 22 tissue types simultaneously. This RNA seq based rapid analysis serves as explanatory data to furnish comparative information for a selected gene. A representative boxplot of pan-cancer analysis is displayed in **Figure 1**.

**Figure 1.**
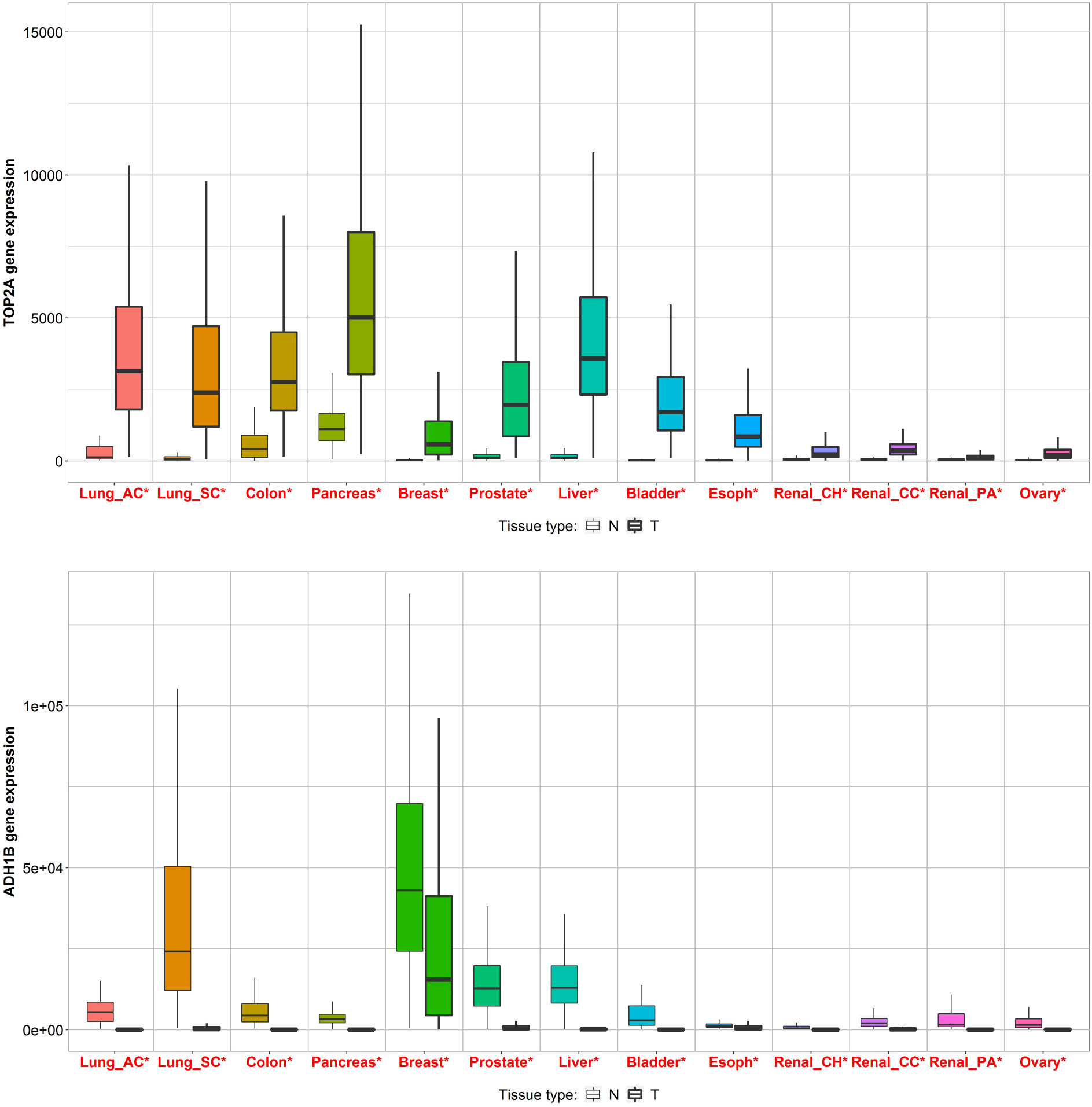
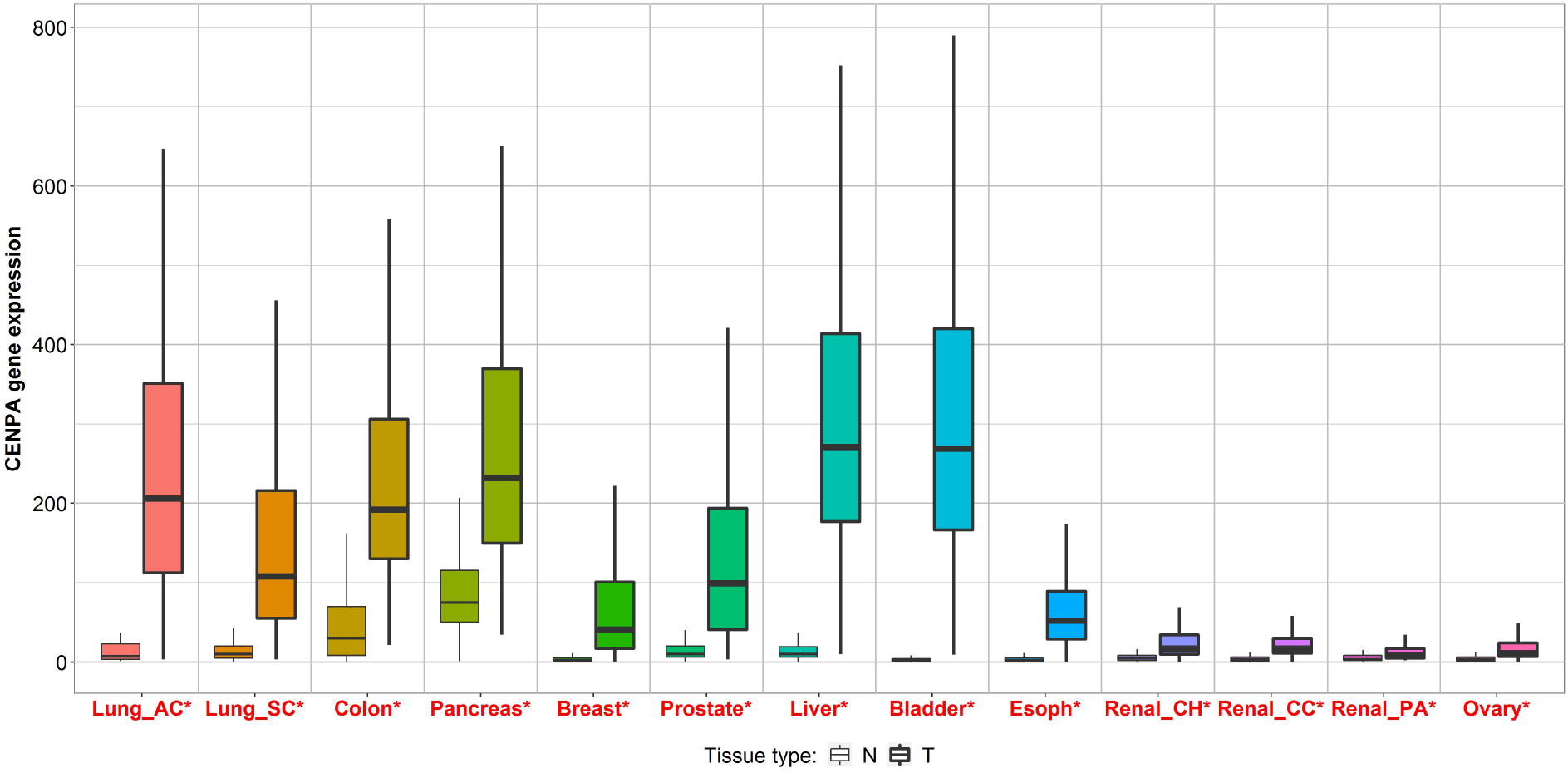
Boxplot of top three genes differentially expressed in most of the ten most common tumor types. Differences significant by a Mann-Whitney U test are marked with red color (*p < 0.01).

The second approach compares directly tumor and normal samples by either grouping all specimens of the same category and running a Mann-Whitney U test or – in case of availability of paired normal and adjacent tumor – by running a paired Wilcoxon statistical test. The results are visualized by both boxplots and violin plots. We have also implemented a graphical representation of sensitivity and specificity: a diagram provides the percentage of tumor samples that show higher expression of the selected gene than normal samples at each major cutoff value. Example outputs of normal-tumor comparison are displayed in **Figures 2 and 3**.

**Figure 2.**
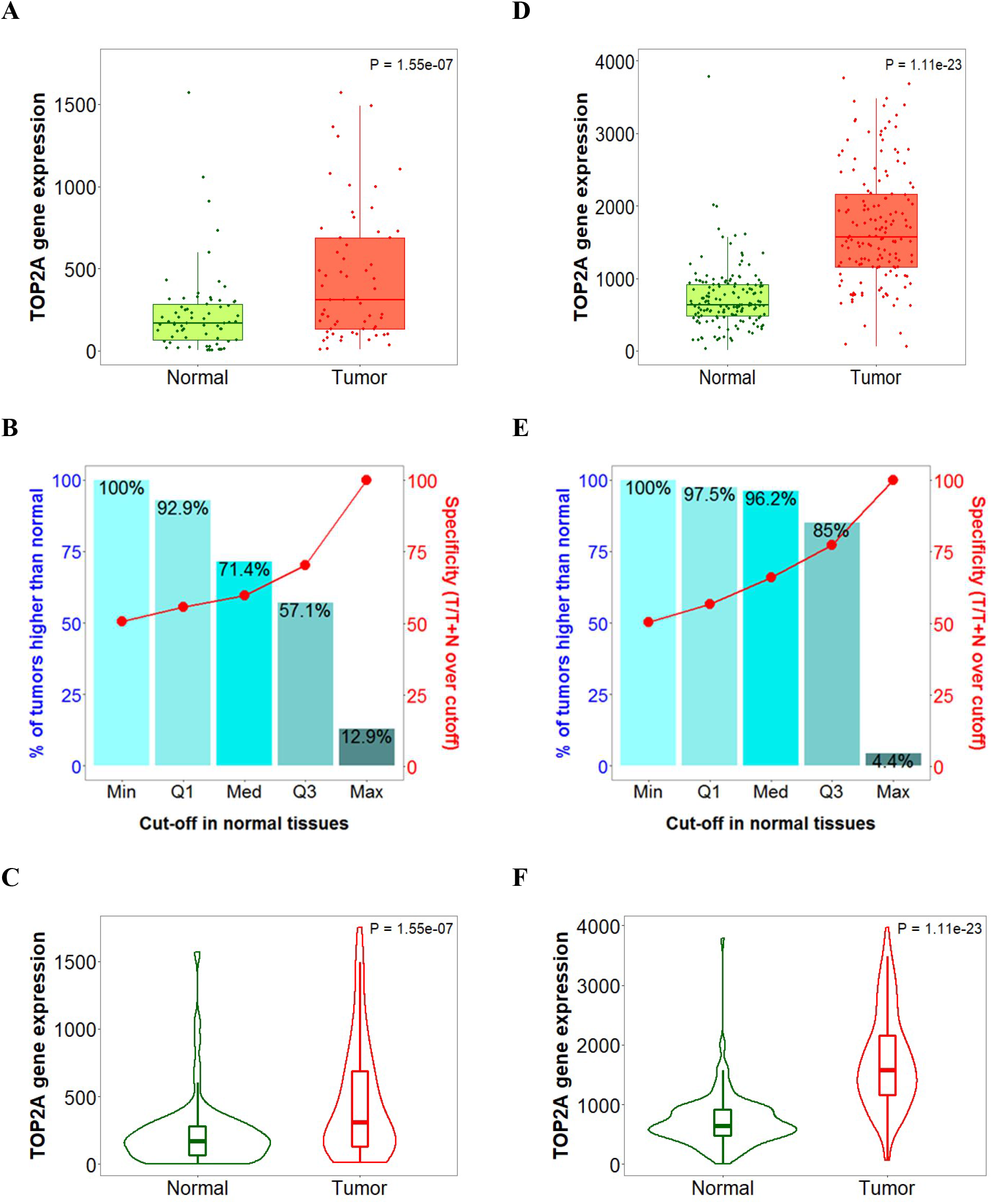
Boxplots (A,D), bar charts (B,E) and violin plots (C,F) of TOP2A gene expression in breast (left) and colon cancer (right) when comparing paired normal and tumor gene array data.

**Figure 3.**
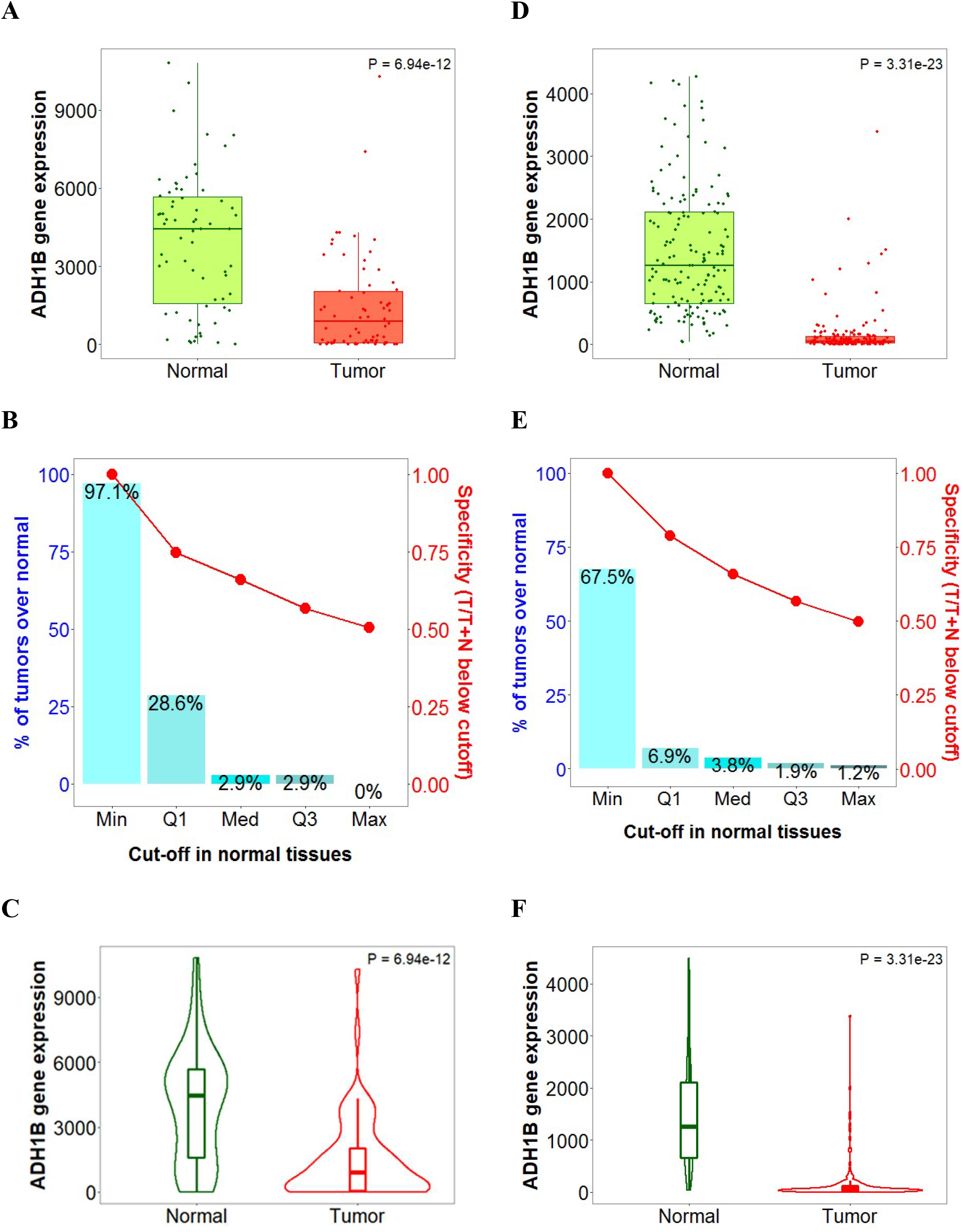
Boxplots (A,D), bar charts (B,E) and violin plots (C,F) of ADH1B gene expression in breast (left) and colon cancer (right) when comparing paired normal and tumor gene array data.

Although the number of metastatic samples is limited in most cases, five and twelve tissue types in the RNA seq and gene array databases have useful amount of specimens. The third feature of the analysis platform allows us to simultaneously compare these tumor, normal and metastatic data using a Kruskal-Wallis test.

### Gene expression analysis of cancers with the highest mortality

We compared the expression of all genes in normal and tumor samples across the ten most lethal tumor types including breast, bladder, colon, lung, liver, esophageal, prostate, pancreas, renal, and ovarian cancer. In the gene array dataset 555 - 2,623 reached statistical significance at FDR <10% and fold change over 1.5. The entire list of all genes is presented in **Supplemental Table 2**. When using the RNA seq cohort, 3,189-12,037 genes were dysregulated at FDR <10% and fold change over 1.5, the entire list of all genes dysregulated in the RNA seq cohorts is presented in **Supplemental Table 3**.

### Linking the most significant genes to cancer hallmarks

We linked the best 55 genes common across all cancer types in both platforms to the cancer hallmarks based on their functions available in Entrez Gene Summary, GeneCards Summary, and UniProtKB/Swiss-Prot Summary. The majority of the genes (n = 21) were linked to sustained proliferative signaling. The second most common hallmark was the deregulation of cellular energetics (n = 13). Activation of invasion and metastasis (n = 5), enabling replicative immortality (n = 8), and avoiding immune destruction (n = 5) were also represented by multiple genes. Only single genes were linked to genome instability and mutation, evasion of growth suppressors, and tumor promoting inflammation. The overlapping 55 genes are listed in **Table 2**.

**Table 2.** Summary of sample numbers in each dataset.

| **Tissue** | **Sample #** | **T** | **N** | **M** | **GEO** | **GTE<sub>x</sub>** | **TCGA** | **TARGET** |
| --- | --- | --- | --- | --- | --- | --- | --- | --- |
| adrenal gland | 531 | 321 | 208 | 2 | x | x | x | - |
| airway epithelia | 99 | 0 | 99 | 0 | x | - | - | - |
| bile duct | 80 | 68 | 10 | 2 | x | - | x | - |
| bladder | 595 | 555 | 40 | 0 | x | x | x | - |
| blood | 407 | 0 | 407 | 0 | - | x | - | - |
| bone | 123 | 123 | 0 | 0 | x | - | - | x |
| breast | 9400 | 8666 | 645 | 89 | x | x | x | - |
| cervix | 565 | 493 | 70 | 2 | x | x | x | - |
| CNS | 4840 | 2825 | 2015 | 0 | x | x | x | - |
| colon | 3120 | 2085 | 935 | 100 | x | x | x | - |
| endometrium | 70 | 39 | 31 | 0 | x | - | - | - |
| eye | 191 | 191 | 0 | 0 | x | - | x | - |
| gastric | 2250 | 1596 | 654 | 0 | x | x | x | - |
| germ cell | 442 | 183 | 259 | 0 | x | x | x | - |
| head and neck | 650 | 599 | 49 | 2 | x | - | x | - |
| heart | 600 | 0 | 600 | 0 | - | x | - | - |
| intestine | 223 | 10 | 210 | 3 | x | x | - | - |
| kidney | 1966 | 1458 | 450 | 58 | x | x | x | x |
| liver | 1805 | 1177 | 604 | 24 | x | x | x | - |
| lung | 3824 | 2890 | 926 | 8 | x | x | x | - |
| lymphoid | 6377 | 6184 | 193 | 0 | x | x | x | x |
| mesothelioma | 139 | 139 | 0 | 0 | x | - | x | - |
| myeloid | 3931 | 3866 | 65 | 0 | x | - | x | x |
| nasopharyngeal | 69 | 56 | 13 | 0 | x | - | - | - |
| neural | 819 | 394 | 425 | 0 | x | x | - | x |
| esophageal | 1750 | 601 | 1120 | 29 | x | x | x | - |
| oral cavity | 61 | 38 | 18 | 5 | x | - | - | - |
| ovarian | 1341 | 1118 | 179 | 44 | x | x | x | - |
| pancreas | 803 | 425 | 360 | 18 | x | x | x | - |
| pancreas -<br>neuroendocrine | 47 | 47 | 0 | 0 | x | - | - | - |
| Parathyroid | 2 | 1 | 1 | 0 | x | - | - | - |
| pituitary | 219 | 18 | 201 | 0 | x | x | - | - |
| prostate | 1098 | 781 | 310 | 7 | x | x | x | - |
| salivary gland | 114 | 10 | 104 | 0 | x | x | - | - |
| skin | 1835 | 356 | 1035 | 444 | x | x | x | - |
| soft tissue | 2878 | 1464 | 1411 | 3 | x | x | x | x |
| spleen | 162 | 0 | 162 | 0 | - | x | - | - |
| thymus | 121 | 119 | 2 | 0 | - | - | x | - |
| thyroid | 1282 | 717 | 557 | 8 | x | x | x | - |
| tongue | 68 | 57 | 11 | 0 | x | - | - | - |
| uterus | 977 | 757 | 220 | 0 | x | x | x | - |
| blood vessel | 913 | 0 | 913 | 0 | - | x | - | - |
| vulva | 151 | 15 | 136 | 0 | x | x | - | - |

**Table 3.**
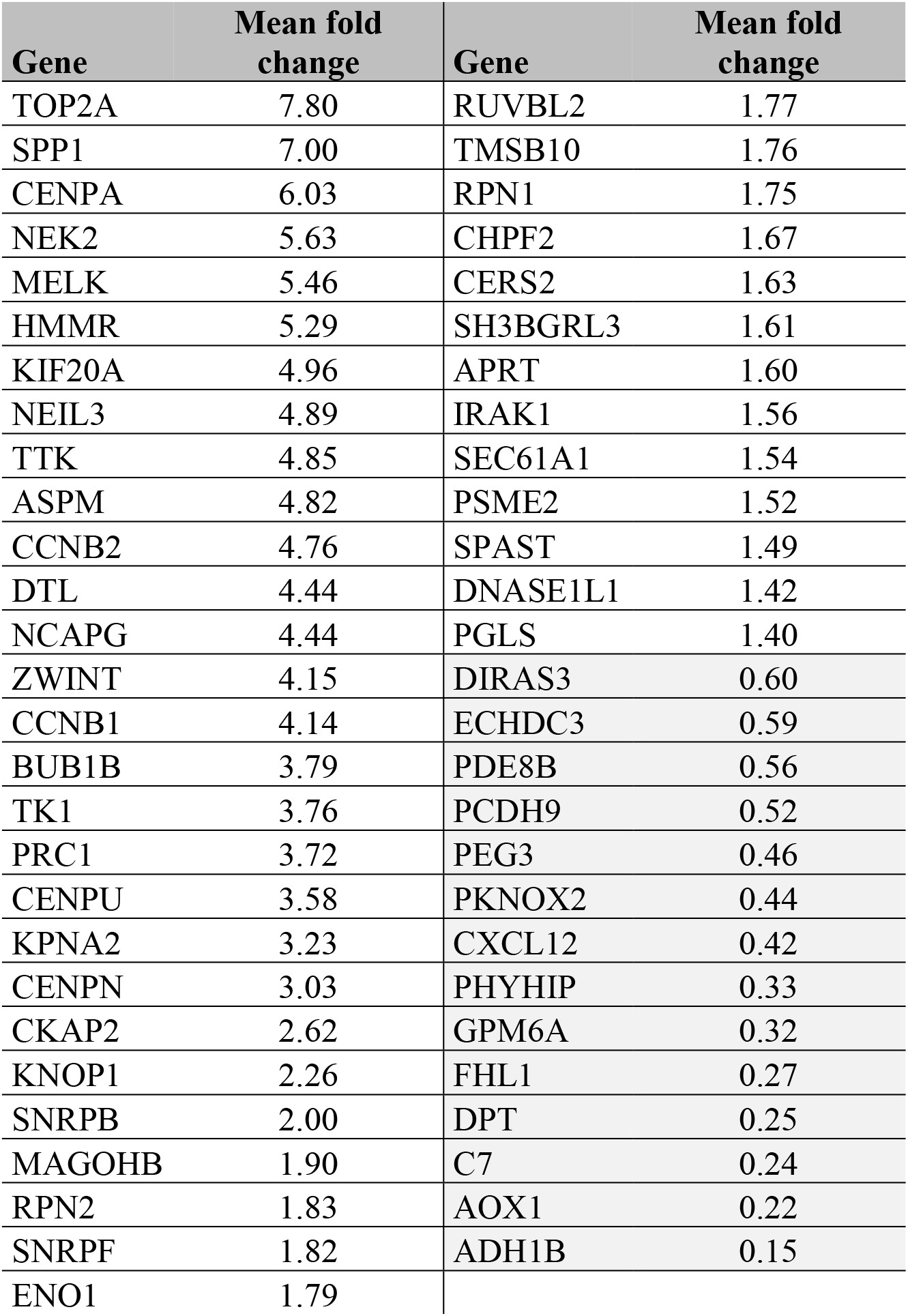
Top fifty-five genes differentially expressed when comparing normal and tumor samples across the ten most common tumor types in RNA-seq and gene array datasets. *Fold change over one corresponds to higher expression in tumors, and fold change below one corresponds to higher expression in normal specimens.*

### Sensitivity and specificity

Whenever a new biomarker is developed, the two most crucial information include sensitivity (the proportion of tumors who have higher expression than normal at a given cutoff) and specificity (the proportion of tumors divided by the total sum of all tumors and normal over the given cutoff). The online analysis interface provides a graphical representation of sensitivity and specificity at the major cutoff values (minimum, Q1, median, Q3, and maximum).

TOP2A was the most upregulated gene in the above analysis with a fold change of 3.26 in breast cancer and 2.54 in colon cancer among others. In **Figure 2**, the expression boxplot, the sensitivity/specificity plot, and the violin plots for TOP2A are displayed using the breast and colon cancer datasets. The most downregulated genes was ADH1B, which had a fold change of 0.22 in breast cancer and 0.3 in colon cancer (see detailed plots in **Figure 3**).

### Validation of differential expression between normal and tumor samples

In order to confirm reproducibility of differential expression and to confirm reliability of the integrated database we conducted a validation using randomly selected training and test cohorts across breast, lung and colon cancers using both RNA-Seq and gene array samples. In each setting, the training and test sets were equally sized to avoid false positive or false negative findings. In the breast cancer gene array and RNA seq datasets all in all 7,223 and 11,689 genes were significant in both training and test sets. These deliver a high concordance in both cases with a chi-square test p value < 0.0001. Regarding colon cancer, 8,259 and 6,763 genes were significant in both training and test dataset in gene array and in RNA seq samples, respectively (p<0.0001). In lung cancer, altogether 7,846 and 8,484 overlapping genes reached significance in both examined cohorts in gene array platform and in RNA seq, respectively (p<0.0001). As each executed analysis showed a p < 0.0001, we conclude that the database can provide highly reproducible results in both platforms. Volcano plots and Venn diagrams depicting results of the validation are listed in **Figure 4**.

**Figure 4.**
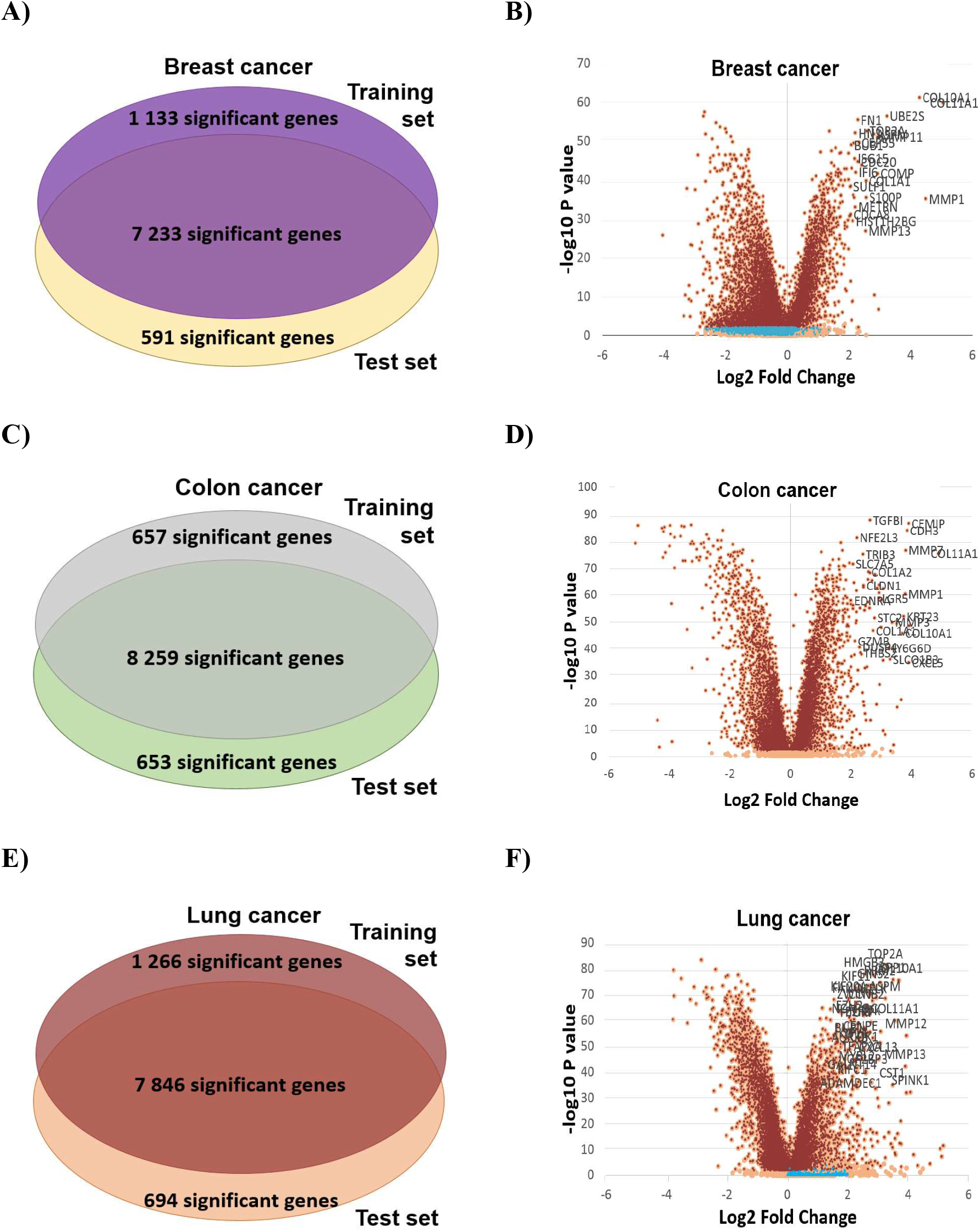
Volcano plots and Venn diagrams of differentially expressed genes in breast, colon and lung cancer using equally sized training-test sets. Venn diagram (A) and Volcano plot (B) from breast cancer. Venn diagram (C) and Volcano plot (D) from colon cancer. Venn diagram (E) and Volcano plot (F) from lung cancer.

## DISCUSSION

Our most important aim was to establish a framework for the comparison of gene expression in malignant, normal and metastatic tissues. To that end, we established a database from publicly available RNA-seq and gene array resources. Followed by a multistep manual and computational curation, we used the datasets in combination with established statistical algorithms to set up an online analysis platform. Finally, the reproducibility of the results delivered by our approach was validated using a training-test approach with multiple randomly differentiated cohorts in two distinct tumor types. Since all implemented examinations delivered high concordance we can state that the established database provides solid results in both platforms used.

One of the major features of our approach is the generation of an expression-cutoff based sensitivity/specificity plot. This graphical representation displays a bar graph showing the proportion of tumor samples with elevated expression compared to the normal cohort at selected cut-off values (minimum, first quartile, median, third quartile, maximum). Since useful pharmacologically targets have to be as specific for the tumor cell as possible, by looking on the graph one can get easily interpretable information regarding the clinical utility of the selected gene. The conventional approach to show sensitivity and specificity would be to generate a receiver operating characteristics (ROC) plot and examine the area under the curve to assess the usefulness of a potential biomarker. Of note, we have recently established the www.rocplot.org platform capable of identifying predictive biomarkers in multiple tumor types by employing ROC analysis [18]. However, one has to set a clinically applied cutoff, thus the overall performance of a marker in a ROC analysis is of little clinical value. Another minor drawback of the ROC plot is that the determination of the optimal cutoff value needs additional computations.

After completing the entire database, our paramount question was: which genes are most specific to cancer across multiple tumor types? We performed a comparative study across the top ten most deadly tumor types and ranked the common genes in these malignancies regardless of the platform. The most consistently upregulated gene was DNA topoisomerase 2-alpha (TOP2A), a gene playing an important role in transcription and replication. Several studies highlighted the importance of TOP2A, and elevated TOP2A expression can serve as a prognostic biomarker in multiple malignancies including lung [19], colon [20], and breast cancer [21]. At present, multiple drugs including doxorubicin, epirubicin or etoposide are widely used in clinical practice to target TOP2A or other topoisomerase gene products [22]. These agents are now used in multiple tumor types including breast cancer [23], leukemias and lymphomas [24, 25].

The most consistently downregulated gene across the investigated tumor types was Alcohol dehydrogenase 1B (ADH1B), a member of the alcohol dehydrogenase enzyme subgroup which serves as an important member in the ethanol, retinol and further alcoholic substance metabolization processes. In concordance with our results, earlier studies came to a comparable conclusion as down-regulation of ADH1B might have a role in multiple cancers, including colon [26], lung [27] or head and neck cancer [28].

A notable limitation of our study is the low number of available metastatic tissues. Although the total number (n=848) seems useful, these represent only 1.5% of the included specimens. Unfortunately, this is an open issue not dealt with in any of the large-scale data collection projects. Another limitation of our database is the lack of data on gene regulation including alternative splicing. Alternative splicing can result in different proteins with dissimilar functions. A future employment of a multi-omic approach in conjunction with the utilization of proteomic data might help to circumvent these issues [29].

In summary, we established the largest currently available transcriptomic cancer database consisting of 57 thousand samples by utilizing multiple RNA-Seq and microarray datasets. We show that the results obtained by these specimens is highly reproducible and have set up a registration-free online analysis portal which enables mining of the database for any gene to assess expression differences in normal, cancer and metastatic samples.

## ACKNOWLEDGEMENTS

The research was financed by the 2018-2.1.17-TET-KR-00001 and KH-129581 grants and by the Higher Education Institutional Excellence Programme of the Ministry for Innovation and Technology in Hungary, within the framework of the Bionic thematic programme of the Semmelweis University.

